# ApproXON: Heuristic Approximation to the E-Field-Threshold for Deep Brain Stimulation Volume-of-Tissue-Activated Estimation

**DOI:** 10.1101/863613

**Authors:** Daniele Proverbio, Andreas Husch

**Affiliations:** University of Luxembourg, Luxembourg Centre for Systems Biomedicine, Interventional Neuroscience Group; University of Luxembourg, Luxembourg Centre for Systems Biomedicine, Systems Control Group; Centre Hospitalier de Luxembourg, National Department of Neurosurgery

**Author notes:** Correspondence (Daniele Proverbio), (Andreas Husch).

## Abstract

This paper introduces a heuristic approximation of the e-field threshold used for volume-of-tissue-activated computation in deep brain stimulation. Pulse width and axon diameter are used as predictors. An open source implementation in MATLAB is provided together with an integration in the open LeadDBS deep brain stimulation research toolbox.

## 1. Background

Visualisation of electrode placement and induced electrical fields is increasingly used in deep brain stimulation (DBS) research and clinic.

Different models have been suggested for the estimation of the volume-of-tissue-activated by the DBS and exact prediction of the volume effected by the stimulation is considered increasingly important for efficient parameter tuning (cf. [1]).

Fully heuristic models capture the complete field modelling and volume-of-tissue-actiavgted estimation process in one approximation. For example the model by Mdler and Coenen [2] is directly estimating the volume-of-tissue-activated using stimulation voltage and impedance as predictors. The model of Dembek et al. [3] also accounts for different stimulation pulse widths and is thus applicable to a wider range of cases.

While the simplified models have clear advantages, like much lower computational costs, limitations include difficulties to account for complex contact configurations as found in multiple-current source simulation using segmented leads. Much more complex finite element methods based models like [4] can be applied in such cases. However, this powerful models predict an electrical field (e-Field) and need additional computation steps to turn the fields into an estimation of the activation volume.

In our experiments we relied on the open SimBio/Fieldtrip based Model described in the supplementary material of [1].

### 1.1. Problem

Currently, openly available models only provide a small set of parameter combinations to compute a stimulation field threshold needed to derive a Volume-of-Tissue-Activate (VTA) from an e-Field. In this paper, we propose a straightforward fit of an approximation to published data on pulse width, axon diameter and resulting e-Field threshold. Open Source Matlab models are provided for convenient use.

## 2. Model

We define a log-linear model of the form

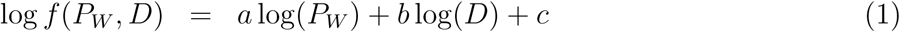

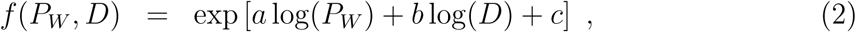

where *P*_*W*_ represent pulse width and *D* axon diameter.

The model was fitted to the data published in Table 3 of Åström et al. [5] in a non-linear least-squares sense using Matlab Curve Fitting Toolbox. This data from [5] is expected to be accurate for a stimulation voltage of 3*V*.

The coefficients *a, b, c* for or model are reported in Table 1. The goodness of fit is estimated by considering a reduced R-square statistics over the degrees of freedom of the fit. In this case, R-square_*red*_ = 0.9948 ∼ 1.

**Table 1:** Model coefficients and confidence bounds.

| Coefficient | Value | 95% confidence bound |
| --- | --- | --- |
| a | -0.6513 | (-0.6819, -0.6206) |
| b | -1.621 | (-1.68, -1.561) |
| c | 3.046 | (2.916, 3.176) |

Figure 2 visualizes the fitted model surface for pulse width *P*_*W*_ ∈ (1, 240) and axon diameter *D* ∈ (1, 8). The isocontour of the proposed *general heuristics* of *T* = 0.2V/mm as suggested in [1] based on [6] is denoted in red.

**Figure 1:**
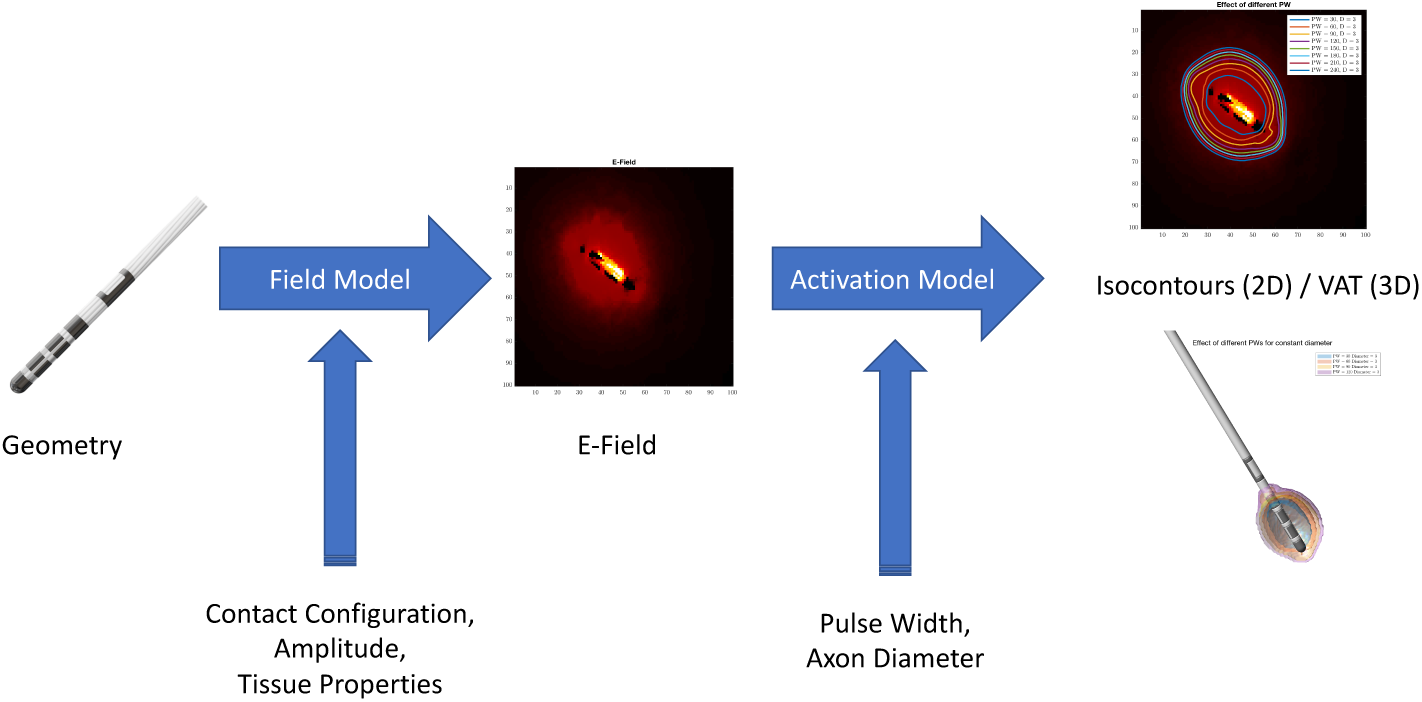
Overview of DBS field modelling and graphical abstract. In this paper, an approximation to the *Activation Model* to threshold an E-Field yielding a Volume-of-Activated-Tissue is presented.

**Figure 2:**
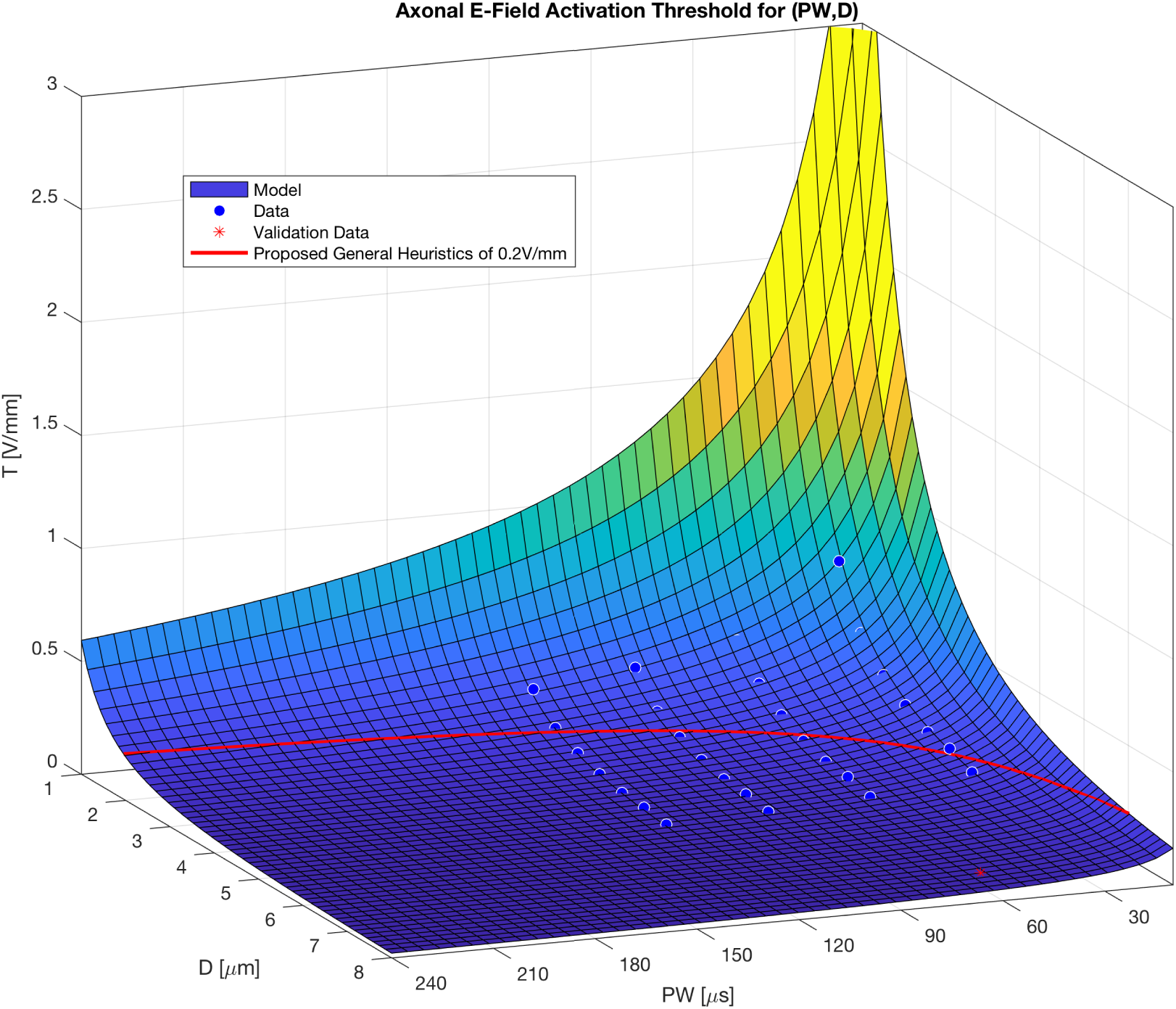
Plot of the model surface predicting the threshold *T* given pulse width *P*_*W*_ and axon diameter *D*. Samples from Table 3 in [5] used for fitting are visualized as circles. An additional point reported in Table 2 in [5] used for validation is denoted by an asterisk. The isocontour of the proposed *general heuristics* of *T* = 0.2V/mm as suggested in [1] is denoted in red.

### 2.1. Derivation of the model structure

Before fitting the parameters of the model to published data, the mechanistic log-linear form of the model was derived as follows.

We first developed a heuristic simplification of the axon electrical and geometrical properties. Considering a heterogeneous manifold of axons in the region around the DBS lead, our minimal model refers to the mean properties of such a manifold and not to the particular geometry or electrical conductance of a single axon. Thanks to this “mean field” perspective, we can thus incorporate heterogeneity in power laws whose exponent are expected to slightly differ from that of a homogeneous population of perfect axons.

Instead of considering a complex geometry as in Åström et al. [5], we applied a geometrical 1th-order approximation, which resembles a cylindrical conducting cable model for the mean axon. In addition, we considered the conductance along the cable as closely ruled by Ohm’s law, after relating ions flow with that of electrons. In this sense, *V*_*T*_ (*P*_*W*_, *D*) is the electric *potential* along the cable. Then *E*_*T*_ = ∇*V*_*T*_ is its gradient, commonly referred as the electric field strength. In turn *f* (*P*_*W*_, *D*) is approximating the threshold for axonal activation under the effect of *E*_*T*_. It will be proportional to the product of *P*_*W*_ (providing energy, cf. [7]) and *D*, that influences the conductance and thus the dampening of electric signal. Because of heterogeneity in shape and electrical properties, their power law dependence has to be fitted with data. Thus, the heuristic model reads:

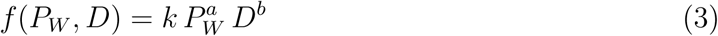

where *f* (*P*_*W*_, *D*) also captures the background potential of the axons manifold, which does not influence a measured difference of potentials. To enable a straightforward fit in the Matlab Curve Fitting toolbox, we then converted the right-hand side of Eq. 3 into an exponential form. By setting log *k* = *c*, Eq. 2 is thus obtained.

## 3. Discussion

Observing Figure 2 it is remarkable that the *general heuristics*, which seems a very reasonable assumption for “typical DBS” parameters in motor-disorders, represents rather extreme axon diameter assumptions for very large respectively very small pulse widths. As the literature indicates that large as well as small pulse width are used more frequently recently, for example large pulse widths in psychiatric indications, the general heuristics assuming a threshold of *T* = 0.2V/mm seems not appropriate for these cases. The presented model allows a more flexible adoption of a threshold given pulse width and fibre-diameter. An example of applying different thresholds computed using the model to an E-Field is shown in Figure 3.

**Figure 3:**
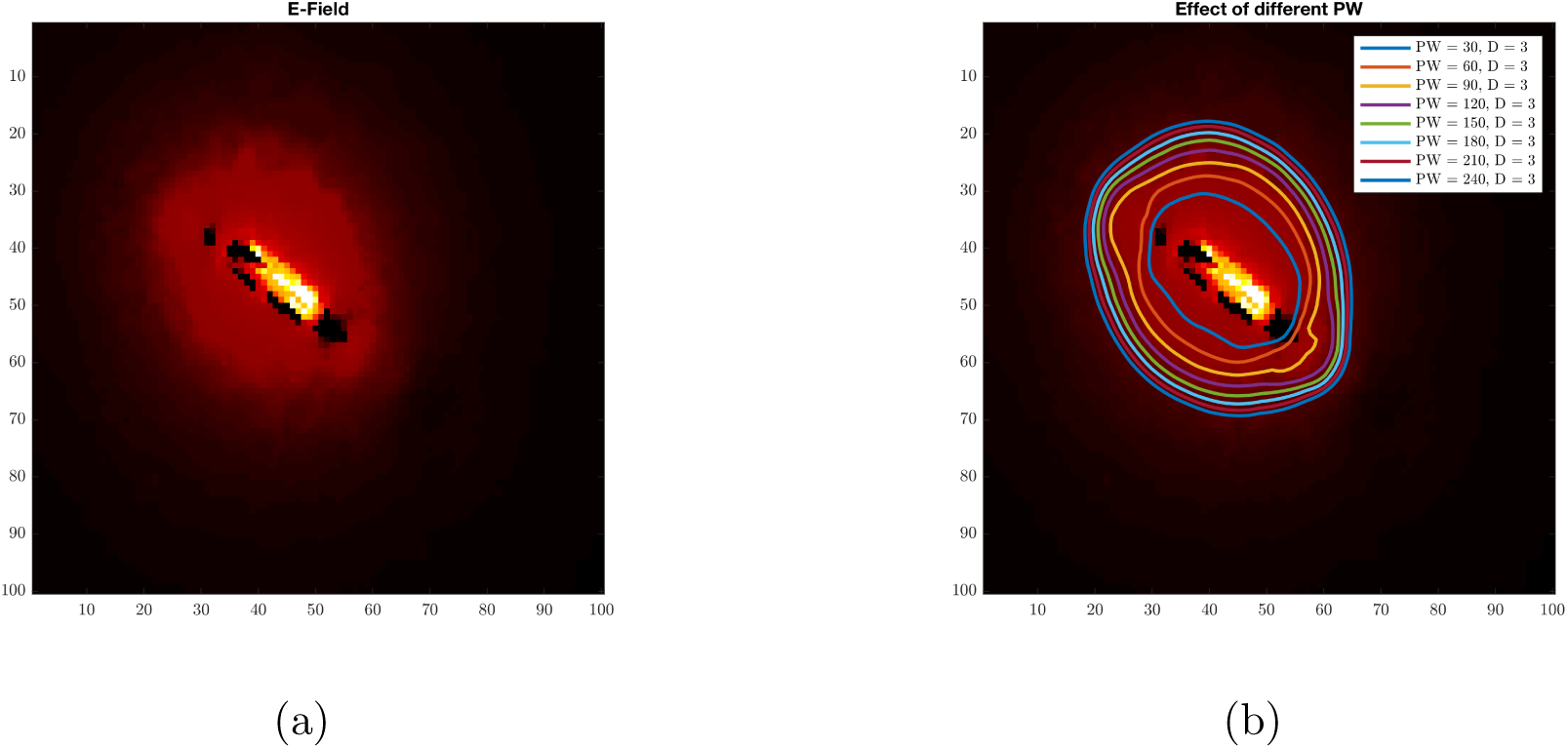
(a) E-Field of a Boston Scientific electrode simulated using SimBio/FieldTrip as implemented in LeadDBS. (b) Overlay of E-Field threshold isocontourlines as predicted by our model for different values of pulse width at a constant axon diameter.

In addition, despite the original data for the fitting being taken at 3*V* [5], the reported influence of the voltage is rather small compared to other factors. Hence, we expect the model to be applicable for other voltages too. In expression we expect that in a realistic setting, for example in post-operative DBS placement analysis, the inaccuracies of other components, like electrode and brain structure detection, are dominating and thus an application of the approximation is still valid.

Figure 4 shows different isosurfaces of an E-Field computed by the model given pulse width and axon diameter pairs.

**Figure 4:**
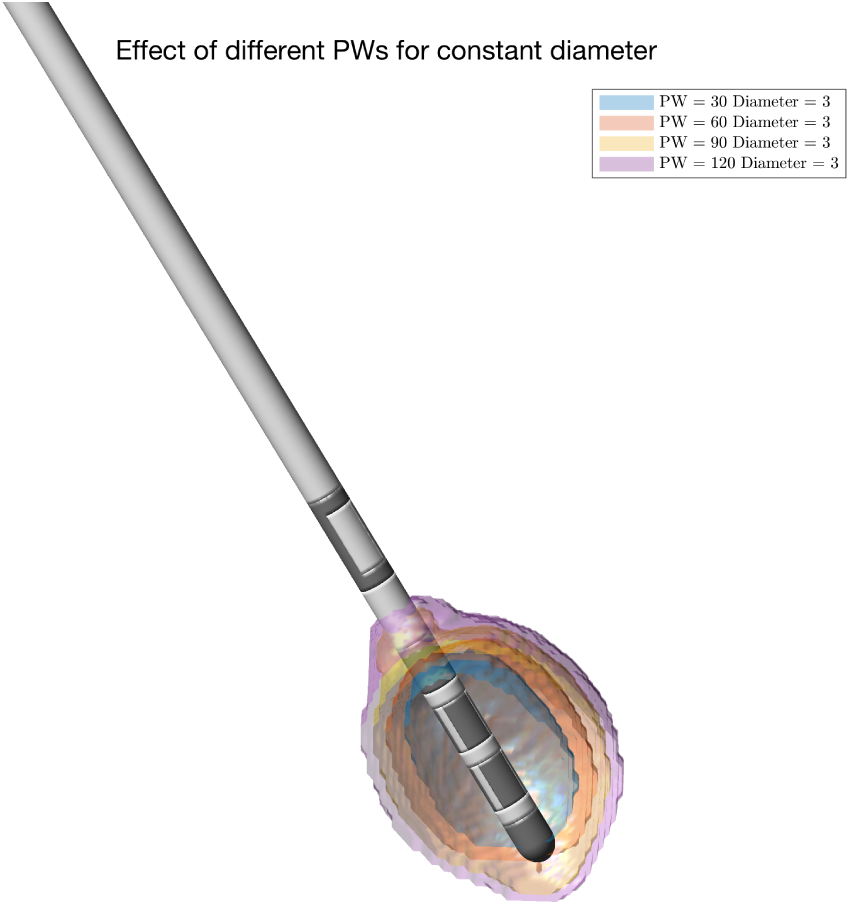
Volume-of-tissue-activated isosurfaces for an E-Field using different thresholds computed by the model given pulse width and axon diameter pairs.

## 4. Conclusion

This report summarized our heuristic approximation used for the published data by Åström [5] on pulse width dependent VAT threshold estimation. We expected it to be very accurate when used for interpolation, use for extrapolation should be done with care. For convenient use open source MATLAB models of the function are provided at https://github.com/adhusch/ApproXON.

For direct use in VAT simulations an open source implementation compatible to the SimBio/FieldTrip DBS Model by Horn and Dembek as available LeadDBS has been contributed to the LeadDBS repository https://github.com/netstim/leaddbs.

Acknowledging the inherent drawbacks of such a straightforward heuristic approach as the presented approximation, the authors hope that future work will enable better models and turn the heuristics superfluous soon. Until that, we hope that the model will have practical use and the released open source code fosters reproducible research.

## Acknowledgement

The Authors would like to thank Prof. Dr. Jorge Goncalves, Systems Control Group, Luxembourg Centre for Systems Biomedicine, University of Luxembourg as well as Prof. Dr. med. Frank Hertel, National Department of Neurosurgery, Centre Hospitalier de Lux-embourg.

